# Affinity maturation of a single-domain antibody via rational mutagenesis informed by artificial intelligence

**DOI:** 10.64898/2026.10.13.758789

**Authors:** Sebastiaan L. Tetteroo, Chelsea Tawiah, Gillian Dekkers, Matti F. Pronker

## Abstract

Antibodies can originate from various sources such as animal immunizations, patient material, DNA/RNA libraries and (AI-based) *de novo* design. Pre-made (*e.g.* synthetic) DNA libraries have substantial advantages over traditional immunization campaigns in terms of speed, cost, animal welfare, amount of required antigen and the possibility to select against otherwise problematic (*e.g.* toxic, unstable or expensive) antigens. The main downside is that selections from these non-immune libraries often result in binders with relatively low affinity due to the absence of *in vivo* affinity maturation. Now that *in silico* structure prediction tools are coming of age, the resulting models could conceivably be used to rationally design mutations, *e.g.* to enhance binding affinity.

Here we used a synthetic single-domain antibody (sdAb) library to select new binders against interleukin-12 (IL-12). One of the selected candidates, 2F3, was predicted to bind to the p40 subunit of IL-12 in a manner that would compete with binding of the receptor protein IL-12Rβ1. Intriguingly, this epitope is also targeted by the clinically approved antibodies Briakinumab and Ustekinumab, which are used to treat a variety of inflammation disorders.

The predicted structure of the IL-12/sdAb complex was used to rationally design eleven point mutations, of which three showed to enhance the binding affinity. By combining these mutations, we could enhance the affinity by eightfold compared to the original lead clone. This shows that correctly (AI-)predicted protein structural models can be used for rational mutagenesis to enhance binding affinity to relevant drug targets.

## Introduction

Single-domain antibodies (sdAbs), such as camelid-derived ‘Variable domain of the Heavy chain of Heavy-chain antibodies’ (VHH, also known as Nanobody®), have many advantages compared to traditional (mammalian) IgG antibodies. They are approximately ten-fold smaller, have greater thermal and enzymatic stability, can be expressed in bacteria and yeast, are easier to chemically conjugate and have better tissue penetration^1,2^. Moreover, they have served as valuable research tools, for example in the fields of protein purification, fluorescence microscopy and structural biology^3–6^, and can resolve preferential orientation problems in cryo-EM^3,5,7,8^. Furthermore, the FDA-approval of Caplacizumab, targeting von Willebrand Factor for the treatment of thrombotic thrombocytopenic purpura, demonstrates that such antibodies have therapeutic potential^9,10^.

Traditionally, sdAbs are obtained through animal immunization followed by phagemid library generation and phage display selection^11^. Selective binders have also been obtained from non-immune libraries^12–14^, which could be either pooled libraries based on previous immunization campaigns, naïve libraries derived from non-immunised animals or synthetic DNA libraries. Advantages of such pre-made libraries include bypassing the process of animal immunization and enabling the use of otherwise problematic (*e.g.* toxic or highly homologous) antigens^15^. However, such libraries lack the *in vivo* affinity maturation by somatic hypermutation combined with selection pressure on affinity, normally occurring in germinal centers of immunised animals. Because of this, sdAbs obtained from non-immune libraries have on average lower affinities compared to those derived from specifically immunised animals^16^. Although such binders can be optimised through various random mutagenesis approaches, new developments in computational structure prediction and *de novo* design of proteins^17,18^ are rapidly reshaping the possibilities to enhance properties such as binding affinity between proteins including sdAbs and their target antigens.

Previously, sdAbs have been successfully developed against various interleukins including Interleukins 12 and 23 (IL-12 and IL-23)^19,20^. IL-12 and IL-23 are secreted heterodimeric cytokines with a shared beta chain called p40, but a differential alpha chain: p35 for IL-12 and p18 for IL-23^21–23^. While they both bind to the receptor IL-12Rβ1 via their p40 subunit, only IL-12 binds to IL-12Rβ2 via p35. In contrast, IL-23 uniquely binds to IL-23R via its p18 subunit^24^. Due to the differential presence of these cell surface receptors in different T-cell subpopulations, IL-12 and IL-23 have distinct signaling outcomes, despite their similarity. IL-12 is a T cell stimulating factor that activates the Th1 pathway leading to differentiation of T cells into Th1 cells, as well as production of interferon-gamma and TNF-α from T cells and natural killer (NK) cells^25^. IL-23 is an inflammatory cytokine that activates the Th17 pathway, resulting in differentiation of T cells into Th17 cells and production of several other (inflammatory) cytokines including Interleukin 17 (IL-17) family cytokines IL-17A and IL-17F^26,27^. Overactivation of the Th17 pathway is involved in many (chronic) inflammation diseases such as Crohn’s disease, ulcerative colitis, plaque psoriasis, psoriatic arthritis, rheumatoid Arthritis, inflammatory bowel disease and multiple sclerosis^28,29^. As such, IL-23 is the target of several widely used therapeutic antibodies such as Briakinumab and Ustekinumab, which both bind to the p40 subunit in a way that competes with IL-12Rβ1 binding, for both IL-12 and IL-23^20,30^.

Here, we selected new IL-12-binding sdAb from a synthetic library by phage display. One of these binders was modelled in complex with IL-12 and predicted to bind to the same site on the p40 subunit of IL-12 as the IL-12Rβ1 receptor. This is the same epitope that is targeted by the clinically approved monoclonal antibodies Briakinumab^20^ and Ustekinumab^30^. We used this model to rationally design eleven point mutations, of which three were confirmed to enhance the binding affinity. By combining these three successful mutations we managed to improve the affinity by more than eightfold, showing that this is a valid approach for affinity maturation.

## Methods

### Phage display

Phage display was performed with the QSyLi synthetic VHH library (QVQ). Phages expressing the VHHs fused to the minor coat protein pIII are based on the VCSM13 bacteriophage and were produced in *E. coli* strain TG1. For production of phages from the library, a total number of bacterial clones that was at least 10 times higher than the number of transformants was inoculated in 2× YT + 2% glucose + 100 μg/ml ampicillin. In case of phage production for subsequent panning rounds, 150 μl of glycerol stocks of TG1 containing rescued phagemids from previous panning rounds were inoculated in 2× YT + 2% glucose + 100 μg/ml ampicillin. The cultures were incubated at 37 °C shaking at 160 rpm until an OD600 of 0.5 was reached and infected with VCSM13 helper phage with multiplicity of infection of >5 phage/bacterium. After 30 min incubation at 37 °C, the cultures were pelleted by centrifugation at 4,200 ×*g* for 12 min and then dissolved in 50 mL 2× YT + 100 μg/ml ampiclin + 25 μg/ml kanamycin. Phages were produced overnight at 37 °C, shaking at 160 rpm. The following day, bacterial pellets were collected by centrifugation at 4,200 ×*g* for 15 min, before adding 40 mL of the supernatant to 10 mL 20% PEG6000/2.5M NaCl, incubating for 1 hour at 4 °C to precipitate phages. Next, phages were pelleted by centrifugation at 4.200 ×*g* for 20 min, supernatant was removed, and phages were resuspended 1 mL phosphate-buffered saline (PBS). Subsequently, bacterial debris was pelleted by centrifugation at 16,200 ×*g* for 5 min, the supernatant was added to 250 μl 20% PEG6000/2.5M NaCl and incubated for 30 min at 4 °C. Following the second PEG incubation, the phages were pelleted by centrifugation at 6,200 ×*g* for 5 min, the supernatant was removed, and the phages were resuspended in 1 mL PBS. One more centrifugation step of 16,200 ×*g* for 5 min was performed to remove the last bacterial cell debris. The phages were stored at 4 °C until use.

Phage display selections were performed on recombinant IL-12 (Sino Biological, CT011-H08H) in three panning rounds. For all three rounds, 500 ng of IL-12 in PBS was coated per well on Nunc MaxiSorp plates (VWR 735-0083, Thermo Fisher Scientific) overnight at 4 °C. The following morning, the wells were washed three times with PBS and blocked with 200 μl 4% of skimmed milk (Marvel) in PBS (MPBS) by shaking for 1 hour. Unless stated otherwise, all shaking steps were performed at room temperature at 900 rpm on a plate shaker (Heidolph Instruments Titramax 100). During blocking of the coated antigens, phages were pre-incubated in 2% MPBS on a rotating wheel at 20 rpm for 30 minutes (IKA Loopster basic). Next, the blocking solution was removed, and 100 µL of phage solution was added and incubated for 2 h while shaking. The phage mix was then removed, and the wells were washed several times with 200 µl PBS with 0.05% Tween20 (PBS-T), followed by washing with 200 μl PBS. Bound phages were eluted with 100 µL of 0.1 M TEA, which was neutralised with 50 µL 1 M Tris/HCl (pH 7.5). Eluted phage outputs were rescued in TG1 and grown overnight with 5 mL 2× YT supplemented with 2% glucose and 100 μg/ml ampicillin at 37 °C shaking at 160 rpm. To assess input and output size of the phages, phages were tenfold serial diluted and rescued by infection in TG1 by 30 min incubation at 37 °C non-shaking. 5 µL of each dilution is spotted on an LB-agar plate with 2% glucose and 100 μg/ml ampicillin and grown overnight at 37 °C non-shaking.

From the output of round three, 92 individual clones were. Periplasmic extracts were produced in 96-deepwell V-bottom plates (Corning, product no. 3960). These were seeded 1:90 in 900 μl production medium and grown for 2.5-3 h. After induction with IPTG, the production was continued overnight at room temperature shaking 800 rpm on an orbital deep-well shaker (Edmund Bühler TiMix 5). The following day, the cultures were centrifuged at 4,000 ×*g* for 10 min, after which the medium was removed, and pellets were dissolved in 120 μl PBS. The samples were frozen at -80 °C for 1 h. Subsequently, bacterial cell debris was removed by centrifugation, and 100 μl of the resulting crude periplasmic extracts were harvested and used for ELISA as indicated below. These 92 clones were further characterised by Sanger sequencing, and their sequences were analysed using the PipeBio Antibody Sequence Analysis platform (Benchling). Sequences were annotated for VHH, using IMGT Alpaca immunoglobulin sets^31,32^. Subsequently, the VHH sequences were clustered for >80% CDR-H3 homology.

### Production and purification of sdAb

To produce the selected sdAb clones, the pPQ81 plasmids containing the genes encoding the candidates were transformed into *E. coli* strain BL21 Rosetta™ 2 (Novagen®, product. no. 71402). The QSY-2F3 mutants were ordered as clonal genes in plasmid pMEK222 with C-terminal FLAG-and His_6_-tag (AAADYKDDDDKGAAHHHHHHGAA; Twist Bioscience) and subsequently transformed into *E. coli* TG1. Cultures were grown overnight in 2× YT supplemented with 2% glucose and 100 μg/ml ampicillin at 37 °C while shaking at 160 rpm. Unless stated otherwise, all following growth and production incubations are incubated at this temperature and speed. Overnight precultures were seeded 1:50 in 50 mL production medium with 0.1% glucose instead and grown for 2.5-3 h until an OD600 of 0.6 is reached, after which antibody expression was induced for 4 h by addition of 1 mM isopropyl-b-D-thiogalactopyranoside (IPTG; Sigma-Aldrich) to the production medium. After production, cultures were pelleted by centrifugation at 3,500 ×*g* for 10 min, dissolved in 3 mL PBS and frozen overnight at -20 °C to lyse the cells and release the periplasmic fraction. The sdAbs were purified from the supernatant using immobilised metal affinity chromatography via the His_6_-tags. The pellets were thawed and spun down, after which the supernatant was incubated 1:20 (v/v) with ROTI®Garose-His/Co Beads (Carl Roth, Art. no. 1235) for 2 h shaking at room temperature in Whatman™ UNIFILTER™ 24-well microplates (Cytiva, product no. 28420850). After incubation, the beads with bound sdAbs were washed subsequently with phosphate-buffered saline (PBS) with 0.05% Tween20 (PBS-T) and PBS. The pre-elution and elution were performed with 15 mM and 150 mM imidazole in PBS, respectively. The eluate was desalted using a AcroPrep™ 24-well 3K Omega filter plate (Cytiva; product no. 97052) in combination with a vacuum manifold. PBS was used for pre-washing the filter, post-washing the sdAbs, and resuspending the sdAbs. Protein concentrations were determined by UV-Vis spectroscopy at 280 nm on a Multiskan^Tm^Go (Thermo Fisher). Protein purity and integrity were checked by sodium dodecyl sulphate polyacrylamide gel electrophoresis (SDS-PAGE).

### Enzyme-linked immunosorbent assays

To detect binding of sdAbs to IL-12, Nunc MaxiSorp plates were coated with 100 ng IL-12-biotin (Acro Biosystems, IL2-H8210) in PBS and incubated overnight at 4 °C. Subsequently the plates were washed three times with PBS-T. All washing steps were performed similarly. Next, plates were blocked with 200 μl 4% MPBS for 1 hour. Unless stated otherwise, all following incubation steps were performed in 50 μl 1% MPBS for 1 hour at room temperature at 900 rpm. After blocking, sdAbs were diluted and incubated on the plate for 1 hour. Purified sdAbs were diluted to a starting concentration of 1 μM and 10 μM and serially diluted 1:10 and 1:5, respectively. sdAb containing crude extracts were diluted 1:5 (10 μl extract with 40 μl 1% MPBS). After the binding of the sdAbs, the plate was washed and incubated with 1:2000 (v/v) diluted primary polyclonal rabbit anti-VHH antibody QE19 (QVQ) for 1 hour. Next, the plate was washed and incubated with 1:5000 (v/v) diluted secondary polyclonal donkey-anti-rabbit-HRP antibodies (Jackson ImmunoResearch). Plates were washed three times with PBS-T and three times with PBS and then incubated with either 7.4 mM (for crude extracts) or 3.7 mM (for purified sdAb) o-phenylenediamine dihydrochloride (OPD, Alfa Aesar) as substrate in 50 mM Na_2_HPO_4_ (Roth) and 25 mM Citric acid (VWR) supplemented with 0.03% H_2_O_2_ (Sigma Aldrich). The reaction was quenched with 25 μl 1 M H_2_SO_4_ and the absorbance was measured at 490 nm on the Multiskan^Tm^Go (Thermo Fisher). Experiments were performed in either singlet or triplicates. Background was determined as the values obtained at the lowest sdAb concentration tested and this was subtracted. Data were fitted according to a one-site, specific binding fit using Graphpad Prism (version 11.1.0 (224)).

### AI-informed rational mutagenesis

The structure of the IL-12-2F3 complex was predicted using Alphafold3^33^. The N-linked glycosylation sites on residues Asn93 and Asn107 of chain α and Asn135 and Asn222 (UNIPROT numbering) of chain β of IL12 were modeled as the common stem of N-linked glycans with two N-acetylglucosamine residues (NAG-NAG). The tryptophan C-linked mannosylation on Trp319 of chain β of IL12 was not modeled. Output models were aligned on the p40 chain in Pymol (Schrödinger) and assessed by standard metrics (pTM score, ipTM score, plDDT, pAE) as well as agreement between the five output models after alignment. The consensus model (that 3 out of the 5 predictions agreed on) was used to inform rational mutagenesis.

### SPR to determine affinity of QSY-2F3 variants to IL12

A White Fox fiber optic surface plasmon resonance (FO-SPR) sensor (Vixen Bio, Carterra) was used to determine the binding affinity of our VHHs to IL-12. All incubation steps were done in PBS, pH 7.4, supplemented with 0.01% Tween-20 at 26 °C, shaking 1000 rpm. 10 nM IL-12-biotin was immobilised for 600 seconds on streptavidin sensor probes (Vixen Bio). After establishing a stable baseline, probes were incubated with increasing concentrations of sdAbs, for 400 seconds per step (kinetic titration). For QSY-2F3_WT a two-fold titration range starting at 4.88 nM until 10.0 µM was used. For the QSY-2F3 mutant a four-fold titration range starting at 244 pM until 1.00 µM was used. Kinetic titration was followed by a dissociation step of 1200 seconds in the buffer. FO-SPR data were collected by FOx software (V2.3.0.139, Vixen Bio), processed using FOx data processing software (V2.0.5.18, Vixen Bio) and on- and off-rates were fitted using TraceDrawer (Version 1.10.1), using a 1:1 model with local *B*_max_, *k*_on_, and *k*_off_ and a constant BI. Equilibrium data was fitted using 1:1 model (Langmuir isotherm). Data was visualised using GraphPad Prism (version 11.1.0 (224)).

## Results

### Selection of IL-12 binders

The selection of IL-12-binding sdAbs was done by three rounds of phage display with the QSyLi synthetic sdAb library. After the third round, the output phage titer from the IL-12 coated wells was substantially higher than that from the non-coated wells. Single clones were picked from this round and sequenced. These clones were also produced as sdAbs and they were subjected to a screening for IL-12 binding in ELISA using crude extracts. This revealed that sdAbs from four different CDR3 clusters showed average IL12-binding signals of more than two times background (Fig. 1A). From each cluster a representative clone was selected, which was produced and purified, except for cluster number 4 which contained a single clone with insufficient sequence quality. Binding of purified sdAbs QSY-2C3, QSY-2F3, and QSY-2G1 from clusters 7, 6, and 9, respectively, to IL-12 was confirmed in a dose response ELISA (Fig. 1B, Table 1). For all three candidates, a full dose response curve up to saturation was not reached within the concentration range tested due to high-concentration assay interference (Hook effect). Nevertheless, out of the three candidates, clone QSY-2F3 appeared most promising with an estimated apparent affinity (*K*_D_) of ∼69 nM, and will be referred to as 2F3 from now onwards.

**Figure 1.**
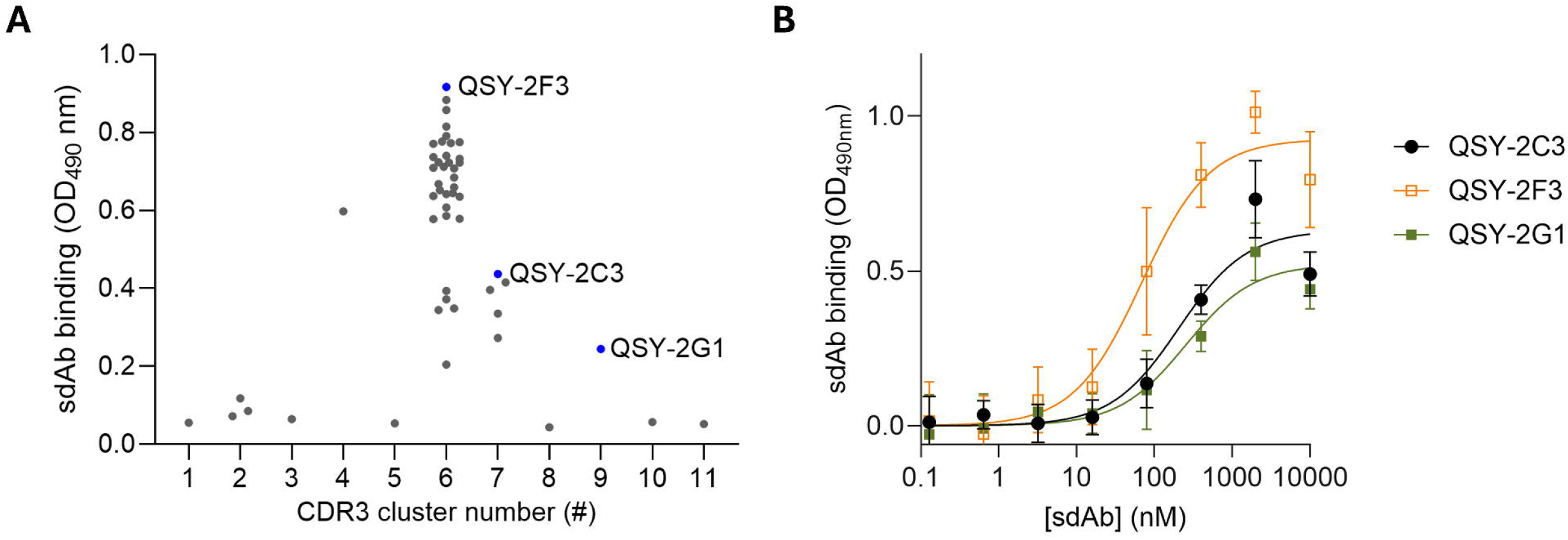
Screening and validation of IL-12 binding sdAbs selected from QSyLi. A) IL-12 binding of CDR3-sequence clustered clones in ELISA with crude extracts shows four CDR3 clusters with IL-12 binding >2 times background (OD_480_ nm of 0.05), selected clones are indicated in orange. Data shown is from a single experiment. B) ELISA dose-response curve of the wildtype QSY-2 clones binding to immobilised IL-12 Bound sdAbs were detected using rabbit-anti-VHH and HRP-conjugated donkey-anti-rabbit antibodies and OPD as substrate. Data represent mean values from a single experiment conducted in technical triplicates.

**Table 1.**
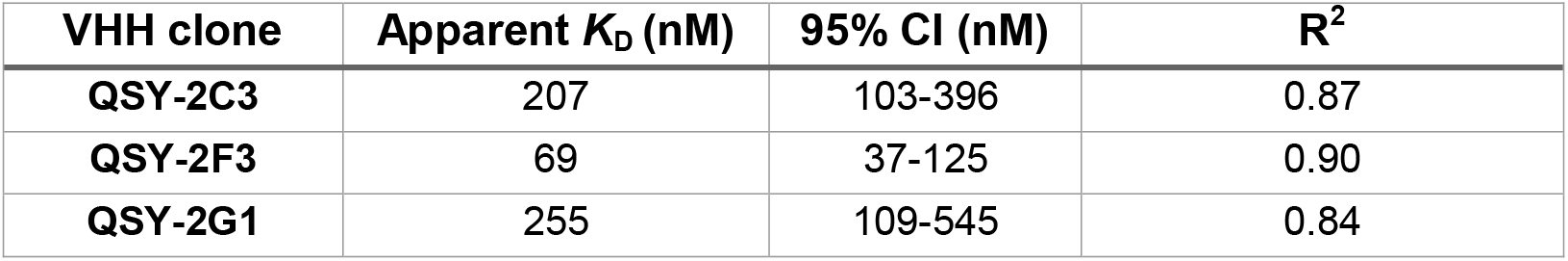
Apparent binding affinities of the selected QSY-2 clones on IL-12. Apparent *K*_D_ values, 95% confidence intervals (CI) and coefficients of determination (R^2^) were derived from nonlinear regression analysis using a one-site specific binding model applied to direct ELISA binding curves (Fig. 1B). Data represent mean values from a single experiment conducted in technical triplicates.

### Rational mutagenesis informed by AI-based structure prediction

To substitute for the *in vivo* affinity maturation normally occurring in the germinal centers of the immunised animal, we sought to improve the affinity of the candidates by using the recent structure prediction method Alphafold3^33^ in combination with rational mutagenesis. The structure of the IL-12 heterodimer in complex with the 2F3 lead VHH sequence was predicted with moderate confidence (ipTM of 0.66, pTM of 0.74, Fig. 2A), with three out of the five predicted structural models converging on a similar binding mode. In these models, 2F3 was found to bind in the wedge between Ig domains 1 and 2 of the IL-12 subunit p40. Interestingly, this is the same epitope as that for the IL12Rβ1, as well as the therapeutic antibodies Ustekinumab^30^ and Briakinumab^20^.

**Figure 2.**
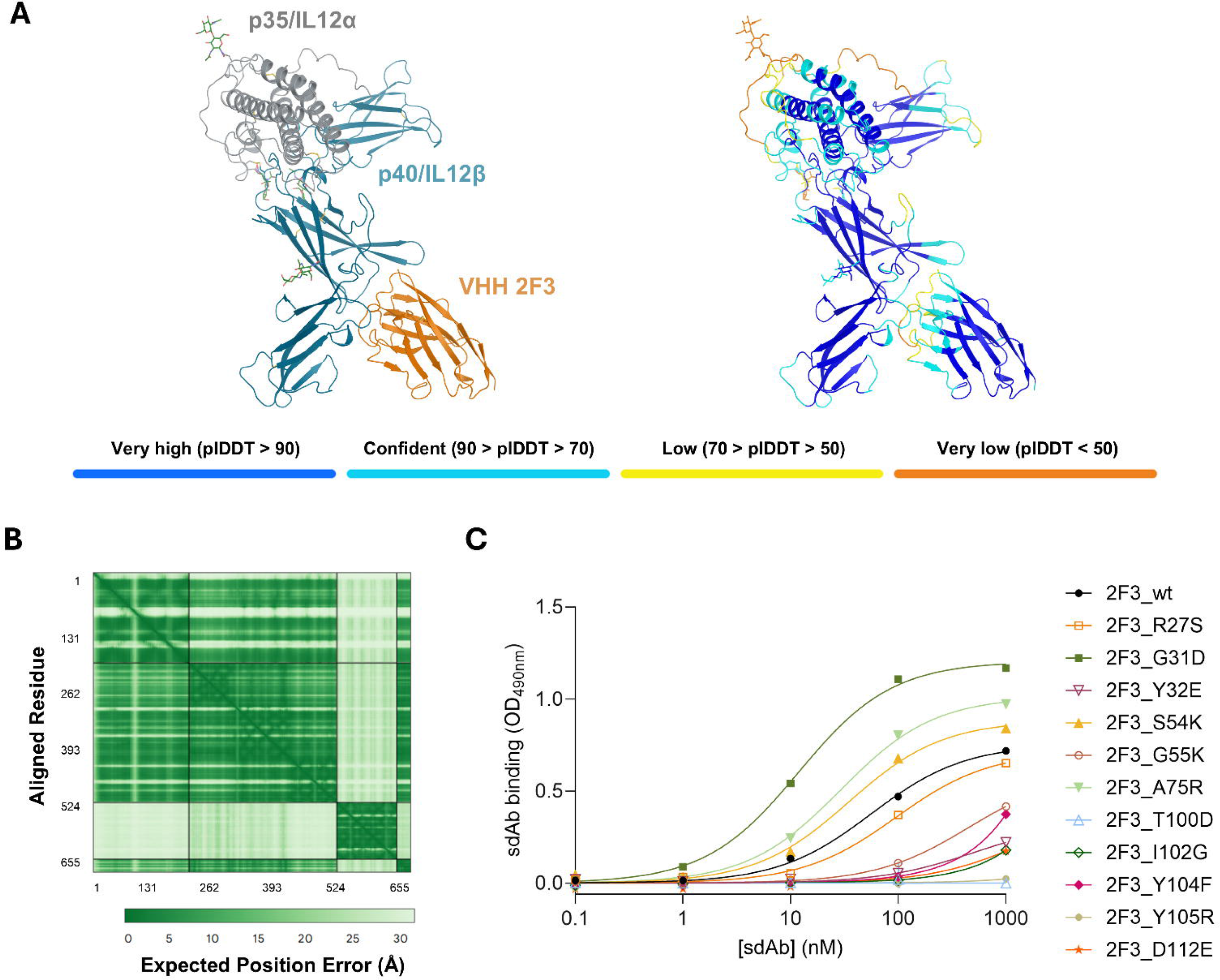
Rational mutagenesis informed by AI-based structure prediction is used successfully for affinity maturation. A) Structure prediction of the IL-12 heterodimer in complex with the lead sdAb 2F3 (left panel) and coloured by plDDT score (right panel). Disulfides and N-linked glycans are shown as sticks. B) predicted Aligned Error (pAE): Expected Position Error versus Aligned Residue C) ELISA dose-response curve of the wildtype QSY-2F3 (2F3_wt) and single amino acid substitution mutants binding to immobilised IL-12. Bound sdAbs were detected using rabbit-anti-VHH and HRP-conjugated donkey-anti-rabbit antibodies and OPD as substrate. Data represent values from a singlet measurement from a single experiment.

The consensus structural model was used as the basis for rational mutagenesis of 2F3. Two structure-informed methods were employed to improve the affinity: reducing putative clashes (R27S, Y104F and I102G) and introducing extra salt bridges for stronger electrostatic interactions (G31D, Y32E, S54K, G55K, A75R, T100D, Y105R and D112E). The mutations were deliberately chosen based on their proximity to the epitope, rather than relying on preconceptions such as CDR loop contributions. These clones were ordered in expression plasmids, produced in *E. coli* and purified. ELISA essays indicated that the strategy of introducing extra salt bridges appeared to be more successful. The G31D mutant showed a 5-fold lower apparent *K*_D_ value compared to the original 2F3 clone (12 nM versus 56 nM, respectively; Fig. 2C, Table 2). The S54K and the A75R mutations either slightly improved or did not affect the *K*_D_ values compared to 2F3 wildtype, indicated that these mutations were tolerated. The remaining eight mutations did not result in lower apparent *K*_D_ values. Notably, for T100D and Y105R, binding was even completely impaired within the tested concentration range. Taken together, the AI-based rational mutagenesis showed potential, as three out of the eleven mutations enhanced the apparent affinity. Moreover, these mutations all originated only from the extra salt bridge introduction strategy.

**Table 2.** Apparent binding affinities of the 2F3_wt and the single mutants for IL-12. Apparent *K*_D_ values, 95% confidence intervals (CI) and coefficients of determination (R^2^) were derived from nonlinear regression analysis using a one-site specific binding model applied to direct ELISA binding curves (Fig. 2C). Data represent values from a singlet measurement from a single experiment. Mutants exhibiting no binding or failing to reach saturation were omitted due to unconstrained curve fitting.

| VHH clone | Apparent $K_D$ (nM) | 95% CI (nM) | $R^2$ |
| --- | --- | --- | --- |
| 2F3_wt | 56 | 43-73 | 0.999 |
| 2F3_R27S | 95 | 64-139 | 0.996 |
| 2F3_G31D | 12 | 9.6-14 | 0.999 |
| 2F3_S54K | 35 | 23-53 | 0.995 |
| 2F3_A75R | 29 | 24-35 | 0.999 |

### Combining successful mutations further enhances the affinity

To determine whether the affinity of 2F3 could be further increased by combining the successful and tolerated individual point mutations G31D, S54K and A75R, the possible double mutants and the corresponding triple mutant were produced in similar fashion to the single mutants. ELISA assays indicated that all combination mutants showed higher apparent binding affinities than the wt and the three single mutants (Fig. 3A, Table 3). The 2F3_G31D_A75R mutant and triple mutant showed the biggest increase in affinity, with virtually identical apparent *K*_D_ values. This suggests that S54K has little to no effect in this combination. However, combining S54K with either G31D or A75R did improve the apparent *K*_D_ values of the latter two substantially.

**Figure 3.**
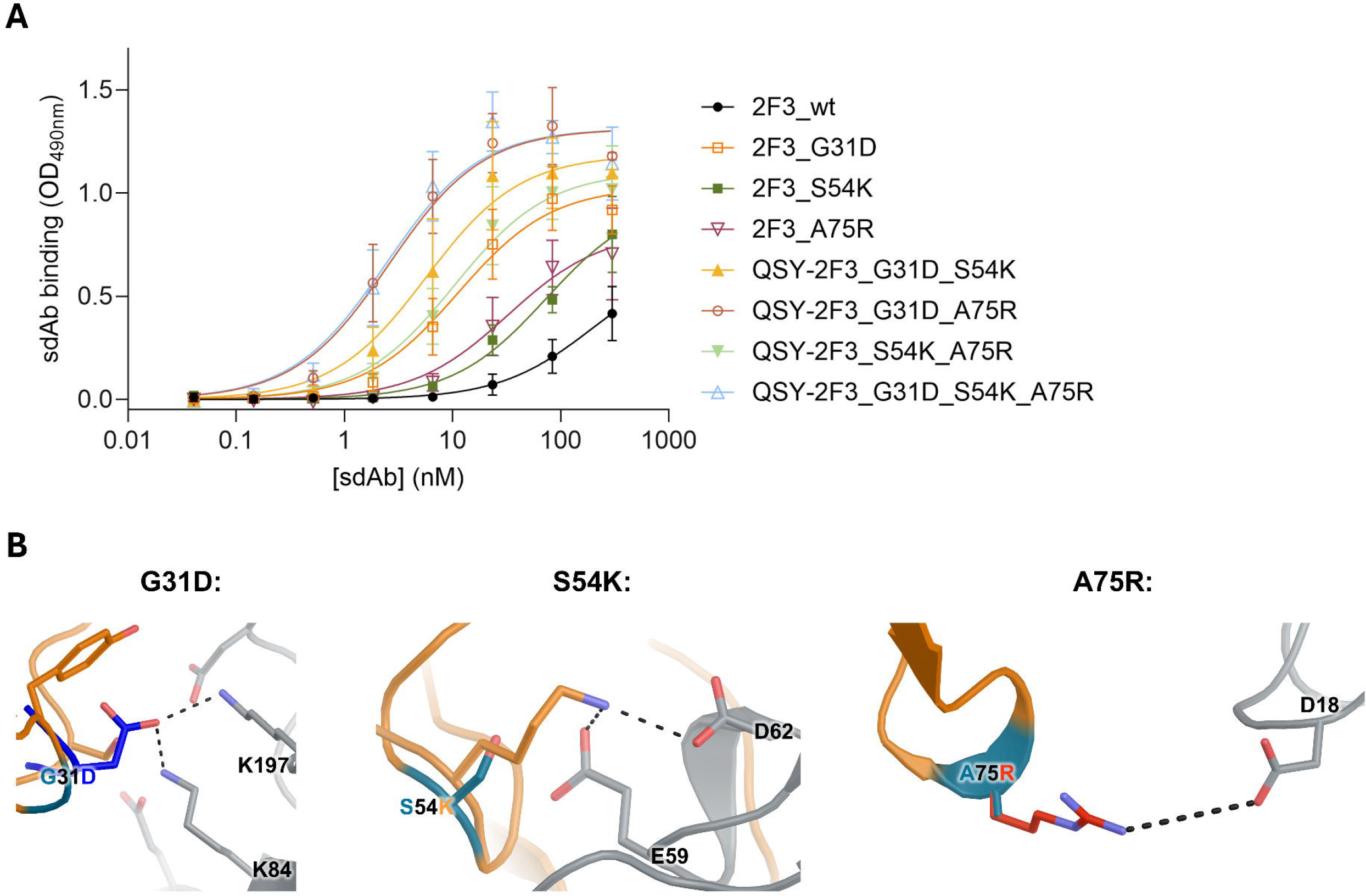
Combining mutations G31D, S54K and A74R successfully increases affinity. A) ELISA dose-response curves of the wildtype 2F3_wt clone, single amino acid substitution and combination mutants binding to immobilised IL-12. Bound sdAbs were detected using rabbit-anti-VHH and HRP-conjugated donkey-anti-rabbit antibodies and OPD as substrate. Data represent mean values from a single experiment conducted in technical triplicates. B) The amino acid substitutions in the G31D (left), S54K (middle) and the A75R (right) mutant introduced extra salt bridges between the sdAb and IL-12, as determined through modeling of the sdAb/IL12 complexes.

**Table 3.** Apparent binding affinities of the 2F3_wt, the successful single, double and triple mutants for IL-12. Apparent *K*_D_ values, 95% confidence intervals (CI) and coefficients of determinations (R^2^) were derived from nonlinear regression analysis using a one-site specific binding model applied to direct ELISA binding curves (Fig. 3A). Data represent mean values from a single experiment conducted in technical triplicates.

| VHH clone | Apparent $K_D$ (nM) | 95% CI (nM) | $R^2$ |
| --- | --- | --- | --- |
| 2F3_wt | 198 | 88-677 | 0.89 |
| 2F3_G31D | 11 | 7.4-17 | 0.93 |
| 2F3_S54K | 74 | 45-126 | 0.95 |
| 2F3_A75R | 32 | 18-58 | 0.92 |
| 2F3_G31D_S54K | 5.7 | 3.6-9.1 | 0.92 |
| 2F3_G31D_A75R | 2.6 | 1.8-3.7 | 0.95 |
| 2F3_S54K_A75R | 10 | 6.7-16 | 0.94 |
| 2F3_G31D_S54K_A75R | 2.4 | 1.6-3.6 | 0.94 |

We also predicted the structure of the triple mutant variant of 2F3 in complex with its target IL12, revealing a similar binding mode as the original (wt) 2F3 clone, but with increased confidence metrics (Fig. 3B). For example, the pTM increased from 0.74 to 0.79 and the ipTM from 0.66 to 0.72. Furthermore, the plDDT and pAE values for contact residues (epitope-paratope) increased substantially. Also, for the triple mutant, five out of five models agreed on the same binding mode as compared to three out of five for the original 2F3-IL12 model. This suggests that, while perhaps not a 1:1 correlation, these confidence metrics might be used as a proxy for binding affinity.

The binding affinities of the sdAbs for IL-12 were quantified by SPR. IL-12 coupled optic probes were subsequently dipped in increasing concentrations of sdAb (kinetic titration) followed by a long dissociation step of 1200 seconds (Fig. 4, Table 4). This showed that the 3 mutations combined affected mainly the association rate constant *k*_a_, with a *k*_a_ of 1.3 × 10^5^ (M × s)^-1^ for the wildtype 2F3 and 7.2 × 10^5^ (M × s)^-1^ for the triple mutant. In contrast, the dissociation constant *k*_d_ was affected only slightly with 2.8 × 10^-2^ s^-1^ and 1.8 × 10^-2^ s^-1^ for the wildtype and triple mutant. Together, these values resulted in a *K*_D_ value of 208 nM for 2F3 wildtype and 25 nM and 2F3_G31D_S54K_A75R, respectively. Equilibrium fits of these data showed very similar results with *K*_D_ values of 200 nM and 25 nM for the wildtype and triple mutant, respectively. Taken together, the ELISA and the SPR experiments showed that the three point mutations increased the affinity of 2F3 by eightfold.

**Figure 4.**
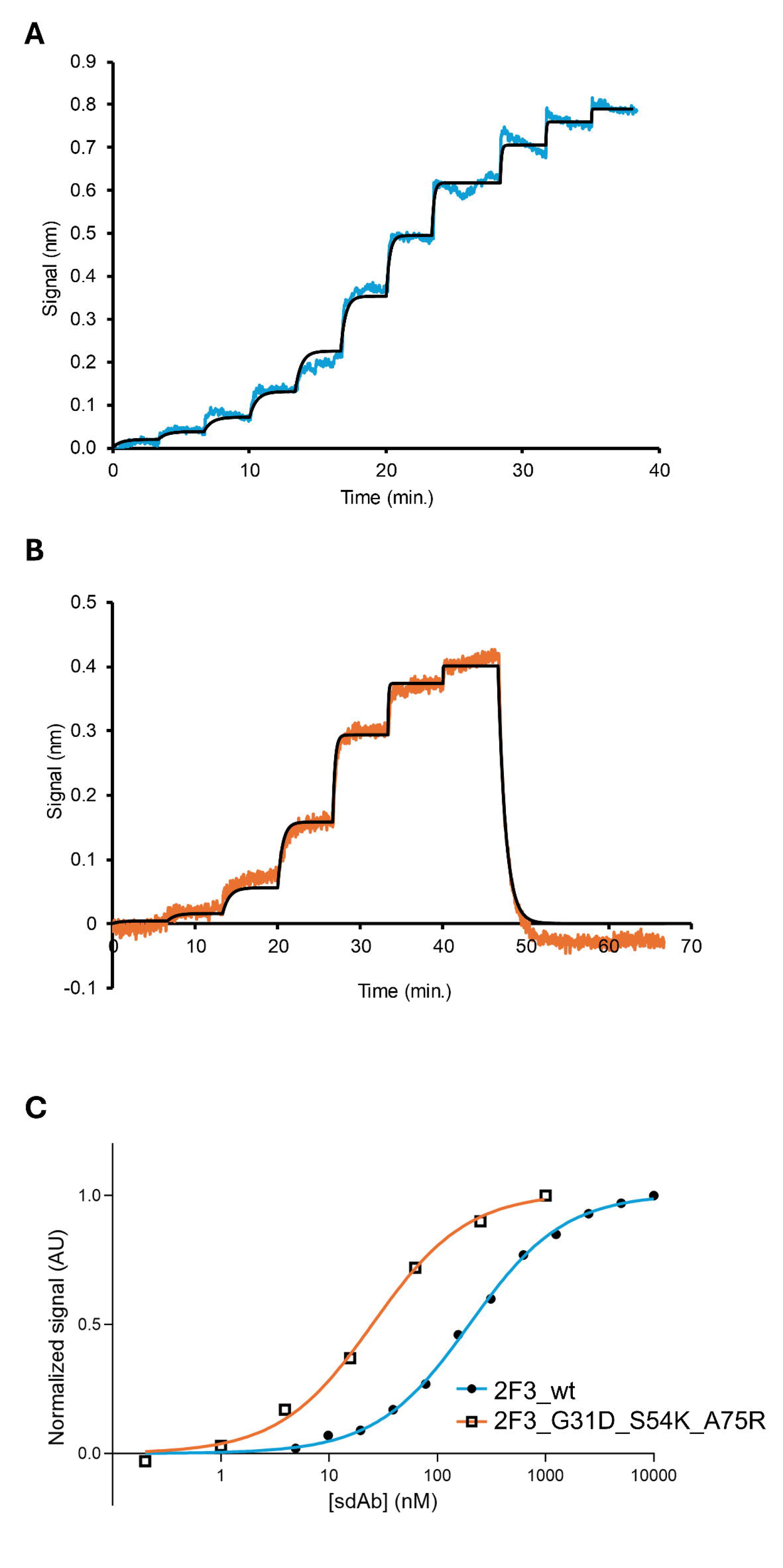
Binding kinetics and affinity of 2F3_wt and 2F3_G31D_S54K_A75R to IL-12 by SPR. A) Kinetic titration of 400 seconds per concentration of 2F3_wt starting at 4.88 nM with two-fold titration until 5 µM. B) Kinetic titration of 400 seconds per concentration of 2F3_ G31D_S54K_A75R starting at 244 pM with four-fold titration until 1 µM. C) Equilibrium fit of both 2F3_wt and 2F3_G31D_S54K_A75R.

**Table 4:** Binding affinities of the 2F3_wt and triple mutant for IL-12 by SPR. Association and dissociation rates from surface plasmon resonance, and binding affinity determined from these rates as well as from an equilibrium fit, in good agreement with each other.

|  |  | *Kinetic titration, 1:1 binding* |  |  | *Equilibrium fit* |
| --- | --- | --- | --- | --- | --- |
| VHH clone | $B_{max}$ (Signal (nm)) | $k_a$ ((M*s)<sup>-1</sup>) | $k_d$ (s<sup>-1</sup>) | $K_D$ (nM) | $K_D$ (nM) |
| 2F3_wt | 0.82 | 1.3x10<sup>5</sup> | 2.8x10<sup>-2</sup> | 208.0 | 200 +/- 7.3 |
| 2F3_G31D_S54K_A75R | 0.41 | 7.2x10<sup>5</sup> | 1.8 x10<sup>-2</sup> | 25.0 | 25.6 +/- 2.5 |

## Discussion

In this study, we describe the selection of novel sdAbs binding IL-12 using a synthetic library by phage display panning, as well as the affinity maturation of the main hit by combining AI-based structure prediction of the antibody-antigen complex, rational mutagenesis and combining successful point mutations. Both the selections and affinity maturation approach were successful at the first try, resulting in a 25 nM affinity sdAb that binds to the same epitope (p40 domain, present in both IL12 and IL23) as the IL12 receptor β1 and as well as the blockbuster drugs Briakinumab and Ustekinumab.

Structure-informed affinity maturation has been used successfully in the past^34^, but many of these efforts precede the rise of AI-based structure prediction tools. Not relying on experimental structure determination is a major benefit in terms of cost and speed, although in terms of accuracy, crystal or cryo-EM structures still are the gold standard. Our affinity maturation approach is in theory more widely applicable and could also be used for conventional antibodies, computationally designed minibinders or VHHs, peptides and natural protein-protein interactions. One caveat is that the approach crucially relies on accurate structure prediction (or experimental structure determination, which would eliminate many of the advantages, such as speed and cost). Furthermore, the program that we used for structure prediction^33^ cannot be used for commercial approaches due to its restrictive license. However, many alternative structure prediction engines without this restriction have been launched since this project was started^35–41^, several of which outperform Alphafold3 on antibody-antigen prediction benchmarks. Moreover, the performance of such structure prediction engines is expected to improve even further in the future. Finally, metrics to assess the quality of these structure predictions are also continuously improving^42–44^.

Since we use the predicted structure as the starting point for our affinity maturation, we were not influenced by preconceptions about the contribution of different structural elements that might be changed to enhance the affinity. For example, the S54K mutation is not in any CDR loop but in the DE-loop, which is also part of the paratope and has previously been suggested to be an overlooked contributor to antibody binding^45^. We also found a mutation in the CDR3 loop (A75R), which has been suggested to be a determinant of specificity rather than affinity^46,47^. G31D also resides in a CDR loop (CDR1), showing that, in contrast to recently suggested approaches for affinity maturation of antibodies^46,47^, CDR loops are solid targets for affinity maturation. Finally, both G31D and S54K are in the middle of the binding interface rather than at the periphery, whereas it has been suggested that mutations at the periphery of this interface are preferred^48,49^. Variation in the Ser52 position (the only residue with its side chain protruding in the DE loop) could potentially be a useful addition in future synthetic library design, as we show here that mutations in this position are well-tolerated and can be beneficial for affinity.

Compared to experimental affinity maturation approaches, such as those relying on error-prone polymerases^50^, mutated oligo pools^51,52^ and combinatorial libraries^53,54^, our approach of testing individual rationally designed point mutations has a major advantage: the successful point mutations can be straightforwardly combined in many cases to further enhance affinity, as we show in this study.

The final frontier would be to avoid the human biases that went into selecting the 11 original point mutations and instead use a fully computational pipeline. However, fully computational methods were recently found to have a success rate not surpassing random hit picking from the original library^55^. While this may change in the future, we find it comforting that human intelligence can still outperform its artificial equivalent for some tasks.

## Author contributions

Conceptualization: GD and MFP. Methodology, investigation and analyses: SLT, CT, and GD for libraries, panning and affinity determinations, and MFP for modelling and affinity maturation. Manuscript writing: MFP, SLT and GD. Reviewing and project management: MFP.

